# Variable Selection in Heterogeneous Datasets: A Truncated-rank Sparse Linear Mixed Model with Applications to Genome-wide Association Studies

**DOI:** 10.1101/228106

**Authors:** Haohan Wang, Bryon Aragam, Eric P. Xing

**Affiliations:** Language Technologies Institute, School of Computer Science Carnegie Mellon University, Pittsburgh, PA, USA; Machine Learning Department, School of Computer Science Carnegie Mellon University, Pittsburgh, PA, USA

## Abstract

A fundamental and important challenge in modern datasets of ever increasing dimensionality is variable selection, which has taken on renewed interest recently due to the growth of biological and medical datasets with complex, non-i.i.d. structures. Naïvely applying classical variable selection methods such as the Lasso to such datasets may lead to a large number of false discoveries. Motivated by genome-wide association studies in genetics, we study the problem of variable selection for datasets arising from multiple subpopulations, when this underlying population structure is unknown to the researcher. We propose a unified framework for sparse variable selection that adaptively corrects for population structure via a low-rank linear mixed model. Most importantly, the proposed method does not require prior knowledge of sample structure in the data and adaptively selects a covariance structure of the correct complexity. Through extensive experiments, we illustrate the effectiveness of this framework over existing methods. Further, we test our method on three different genomic datasets from plants, mice, and human, and discuss the knowledge we discover with our method.

## I. INTRODUCTION

Increasingly, modern datasets are derived from multiple sources such as different experiments, different databases, or different populations. In combining such heterogeneous datasets, one of the most fundamental assumptions in statistics and machine learning is violated: That observations are independent of one another. When a dataset arises from multiple sources, dependencies are introduced between observations from similar batches, regions, populations, etc. As a result, classical methods breakdown and novel procedures that can handle heterogeneous datasets and correlated observations are becoming more and more important.

In this paper, we focus on the important problem of variable selection in non-i.i.d. settings with possibly dependent observations. In addition to the aforementioned complications in analyzing datasets arising from multiple sources, the rapid increase in the dimensionality of data continues to hasten the need for reliable variable selection procedures to reduce this dimensionality. This issue is especially salient in genomics applications in which datasets routinely contain hundreds of thousands of genetic markers coming from different populations. For example, to discover genomic associations for a certain disease, genetic data from patients is often collected from different hospitals. As a result, data from the case and control groups can be confounded with variables such as the hospital, clinical trial, city, or even country. Another common source of sample dependence is family relatedness and population ancestry between individuals [1].

Unfortunately, in many applications information on the origin of different observations is lost either through data compression or experimental necessity. For example, for privacy reasons, it may be necessary to anonymize datasets thereby obfuscating the relationship between different observations. As a result, the data becomes confounded and attempts to learn associations via existing variable selection procedures are doomed to fail [2]. In seeking to discover information from such rich datasets when we do not have this important information, it becomes necessary to *deconfound* our models in order to implicitly account for this.

Existing solutions rely on traditional hypothesis testing after a dedicated confounding correction step, usually resulting in suboptimal performance [3], [4]. In contrast, state-of-the-art variable selection methods usually assume that the data comes from a single distribution, leading to reduced performance when applied to multi-source data.

We directly address the problem of variable selection with heterogeneous data in this paper. Our main contributions are the following:

- We propose a general sparse variable selection framework that takes into account possibly heterogeneous datasets by implicitly correcting for confounders,
- We improve this framework by introducing an adaptive procedure for automatically selecting a low-rank approximation in the linear mixed model,
- We apply our model to three distinct genomic datasets in order to illustrate the effectiveness of the method and report our findings.

## II. RELATED WORK

Variable selection is a fundamental problem in knowledge discovery and has attracted significant attention from the machine learning and statistical communities. The basic idea is to reduce the dimensionality of a large dataset by selecting a subset of representative features without substantial loss of information. This problem has attracted substantial attention in the so-called *high-dimensional* regime, where it is typically assumed that only a small subset of features are relevant to a response. In order to identify this subset, arguably the most popular method is *l*_1_-norm regularization (i.e. *Lasso* regression [5]). More recently, nonconvex regularizers have been introduced to overcome the limitations of Lasso [6]. Examples include the Smoothly Clipped Absolute Deviation (SCAD) [6] and the Minimax Concave Penalty (MCP) [7]. These methods overcome many of the aforementioned limitations at the cost of introducing nonconvexity in the optimization problem; a recent review of these methods can be found in [8]. In applications, variable selection is broadly used to extract variables that are interpretable or potentially causal [9], [10], especially in biology [11] and medicine [12].

When the data is non-i.i.d., such as when it arises from distinct subpopulations, two popular approaches for addressing this are principal component analysis [13] and linear mixed models [2], [14]. Mixed models first rose to prominence in the animal breeding literature, where they were used to correct for kinship and family structure [15]. Interest in these methods has surged recently given improvements that allow their application to human-scale genome data [16], [17], [18], [19], [20], [21]. These methods, however, ultimately rely on classical hypothesis testing procedures for variable selection after confounding correction. Finally, a recent line of work has sought to combine the advantages of linear mixed models with sparse variable selection [22], [23], [24], [25].

## III. TRUNCATED-RANK SPARSE LINEAR MIXED MODEL

Before we introduce our method, we first revisit the classical linear mixed model [26].

### A. Linear Mixed Model

The linear mixed model (LMM) is an extension of the standard linear regression model that explicitly describes the relationship between a response variable and explanatory variables incorporating an extra, random term to account for confounding factors. As a consequence, a mixed-effects model consists of two parts: 1) Fixed effects corresponding to the conventional linear regression covariates, and 2) Random effects that account for confounding factors.

Formally, suppose we have *n* samples, with response variable *y* = (*y*_l_, *y*_2_,…*y_n_*) and known explanatory variables *X* = (*x*_l_,*x*_2_,…*x_n_*). For each *i* = 1,2,…,*n*, we have *x_i_* = (*x_i_*,_l_, *x_i_*,_2_,…*x_i,p_*), i.e., *X* is of the size *n* × *p*. The standard linear regression model asserts *y* = *Xβ* + *∈*, where *β* is an unknown parameter vector and 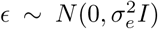. In the linear mixed model, we add a second term *Zμ* to model confounders:

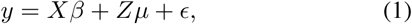

Here, *Z* is a known *n* × *t* matrix of *random effects* and *μ* is a random variable. Intuitively, the product *Zμ* models the covariance between the observations *y_i_*. This can be made explicit by further assuming that 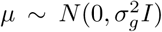, in which case we have

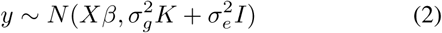

where *K* = *ZZ^T^*. Here, *K* explicitly represents the covariance between the observations (up to measurement error 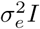): If *K* = 0, then each *y_i_* is uncorrelated with the rest of the observations and we recover the usual linear regression model. When *K* ≠ 0, we have a nontrivial linear mixed model. As K is required to be known, early applications of LMMs also assumed that *K* was known in advance [15]. Unfortunately, in many cases (including genetic studies), this information is not known ahead. In these cases, a common convention is to estimate *K* from the available explanatory variables. As we shall see in following texts, finding a good approximation to *K* is crucial to obtaining good results in variable selection.

### B. Sparsity Regularized Linear Mixed Model

For high-dimensional models with *p* ≫ *n*, it is often of interest to regularize the resulting model to select out important variables and simplify its interpretation. This can easily be achieved by introducing sparsity-inducing priors to the posterior distribution. For example, [24] introduces the Laplace prior, which leads to a *l_l_* regularized linear mixed model as following:

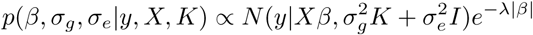

We call the result the *sparse linear mixed model,* or SLMM for short.

This choice of prior—which corresponds to the well-known Lasso when only fixed effects are considered—is well-known to suffer from limitations in variable selection [6], [27]. The first contribution of our paper is to extend this SLMM-Lasso model to more advanced regularization schemes such as the MCP and SCAD, which we call the SLMM-SCAD and SLMM-MCP, respectively. For simplicity, we will use *f* (*β*) to denote a general regularizer, yielding the following general posterior:

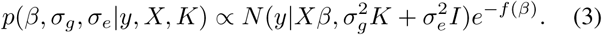

This allows us to combine the (independently) well-studied advantages of the linear mixed model for confounding correction with those of high-dimensional regression for variable selection.

### C. Truncated-rank Sparse Linear Mixed Model

Despite their successes, the main drawback of the aforementioned mixed model approaches is the estimation of *K* from the data *X*. In this section, we propose an adaptive, low-rank approximation for K in order to more accurately model latent population structure as the second contribution of our paper.

#### 1 Motivation

Even though *K* is assumed to be known in LMMs, we have already noted that in practice *K* is often unknown. Thus, to emphasize the distinction between the true, *unknown* covariance *K* and an estimate based on data, we let 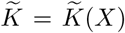 denote such an estimate. Substituting 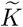 for *K* in (3), the posterior then becomes:

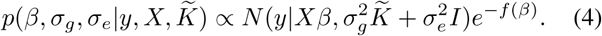

By far the most common approximation used in practice is 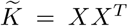 [15], [16]. Under this approximation, equation 1 becomes

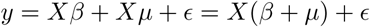

where 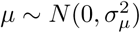. As our goal is the estimation of *β*, this evidently makes distinguishing *β* and *μ* difficult.

This approximation was originally motivated as a way to use the observed variables *X* as a surrogate to model the relationship between the observations *y*. The hope is that the values in *X* might cluster conveniently according to different batches, regions, or populations, which are the presumed sources of confounding. One straightforward observation is that such sources of confounding typically have a much lower dimensionality than the total number of samples in the data. As a result, we expect that *K* will have a low-rank structure which we can and should exploit. Unfortunately, the matrix *XX^T^* will not, in general, be low-rank—in fact, it can be *full rank*, with rank(*XX^T^*) = *n*. To correct for this, we propose the Truncated-rank Sparse Linear Mixed Model (TrSLMM).

#### 2 Method

Instead of choosing 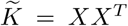 as our approximation, we seek a low-rank approximation to the true covariance *K*. Let Γ:= *XX^T^* and Γ = *UΛV^T^* be the SVD of Γ. Define Λ_*s*_ to be the diagonal matrix such that (Λ_*s*_)_*jj*_ = Λ_*jj*_ for *j* ≤*s* and (Λ_*s*_)_*jj*_ = 0 otherwise (assuming values ofΛ are in decreasing order). Then a natural choice for 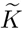 is Γ_*s*_:= *UΛ_*s*_V^T^* for some 0 < *s* < *n*, i.e. the best s-rank approximation to Γ.

*a) Selection of s:* Therefore, we have replaced the problem of estimating *K* with that of estimating an optimal rank *s* from the data. Fortunately, the latter can be done efficiently. To motivate the selection of *s*, we first investigate the distribution of Λ under different population structures. Let *G* denote the number of subpopulations or distributions used to generate the data, which all follows the Gaussian distribution with the zero means. Figure 1 shows a plot of normalized Λ for 100 data samples for *G* = 1; 5; 20; 100. We can clearly see that in the middle two cases (*G* = 5 and *G* = 20), the singular values exhibit some interesting patterns: Instead of decaying smoothly (as for *G* = 1 and *G* = 100), there are a few dominant singular values and more small singular values following a steep drop-off. This confirms our intuition of a latent, approximately low-rank structure within Γ.

**Figure 1:**
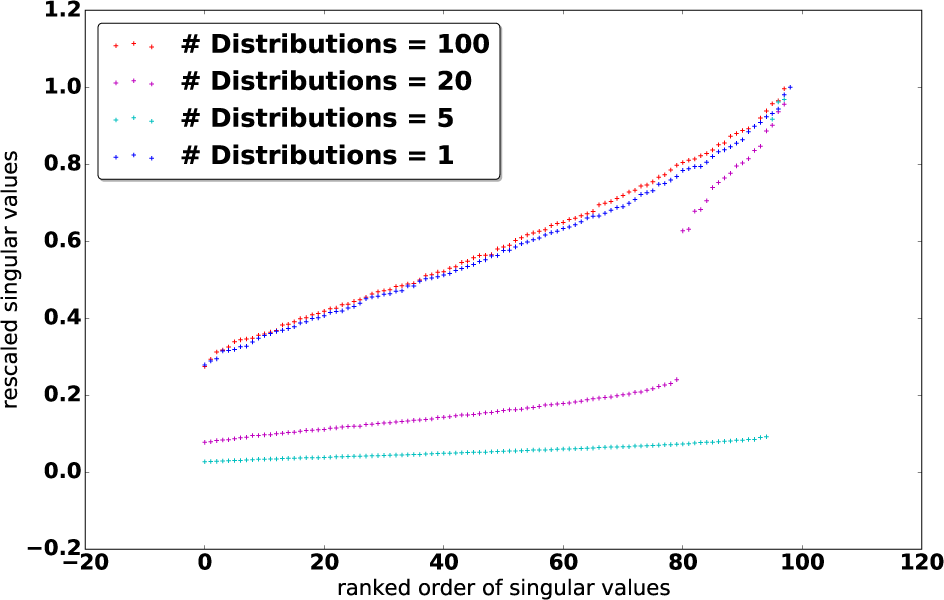
Distributions of singular values of K for different number of distributions the data originate.

Based on this observation, we introduce a clean solution to truncate Λ: We can directly screen out the top, dominant singular values by selecting the top s values Λ _*j*_ for which

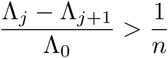

where *n* is the number of samples. In particular, the number of selected singular values *s* satisfies (Λ_*s*_ − Λ_*s*+1_)/Λ_0_ > 1/*n* and (Λ_*s*−1_ − Λ_*s*_)/Λ_0_ ≤ 1/*n*.

Then, we have:

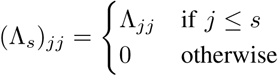

and finally:

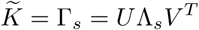

A similar low-rank approximation idea has been used previously [28], [2], however, these procedures require specifying unknown hyperparameters, even when replaced by sparse PCA [29] or Bayesian *K*-means [30]. Another approach is to fit every possible low-rank Λ, sequentially and selecting the best configuration of singular values based on a pre-determined criteria [31], which is *O*(*n*) slower than our method and most importantly does not scale for modern human genome datasets.

#### 3 Parameter Learning

In order to infer the parameters {*β*, *σ*_g_,*σ*_e_}, we break the problem into two steps: 1) Confounder correction, where we solve for *σ*_g_and *σ*_e_; and 2) Sparse variable selection, where we solve for *β* in Equation 4.

##### a Confounder Correction

Following the empirical results in [2], we first estimate the variance term with:

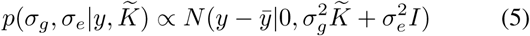

where *y̅* is the empirical mean of *y*. We then solve Equation 5 for *σ*_g_ and *σ*_e_, where we can adopt the trick of introducing 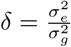 to replace 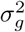 for more efficient optimization [16].

Finally, we can then correct the confounding factors by rotating the original data:

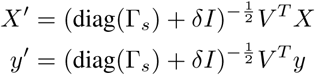

where 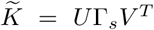 is the singular value decomposition, which has already been computed to determine *s*.

##### b Sparse Variable Selection

After rotating the data to produce *X′* and *y′*, we have a standard variable selection task at hand [24]. Thus, maximizing the posterior in Equation 4 becomes equivalent to solving a variable selection problem with *X′* and *y′*. Note that unlike vanilla linear regression, which would be unchanged by rotations, the introduction of the random effects *Zμ* in (2) violates this rotation-invariance property.

For different choices of regularizer *f* (*β*), we can then solve the following regularized linear regression problem:

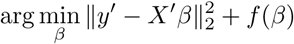

where standard optimization techniques can be adopted. In our experiments, we use proximal gradient descent [32].

## IV. SYNTHETIC EXPERIMENTS

In this section, we evaluate the performance of our proposed method Truncated-rank Sparse Linear Mixed Model (TrSLMM-MCP, TrSLMM-SCAD, TrSLMM-Lasso, as well as SLMM-MCP and SLMM-SCAD) against existing SLMM method (SLMM-Lasso), vanilla sparse variable selection method (Lasso, SCAD, MCP), and recent popular LMM method extensions (LMM-Select [18], LMM-BOLT [20], and LMM-LT [21]).

### A. Data Generation

We first simulate observed covariates coming from *G* different populations. We use *c_g_* to denote the centroid of the gth population, *g* = 1,…, *G*. First, we generate the centroids *c_g_* and from each centroid, we generate explanatory variables from a multivariate Gaussian distribution as follows:

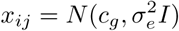

where *x_ij_* denotes the *i*^th^ data from *g*^th^ distribution.

We then generate an intermediate response *r* from *X* from the usual linear regression model:

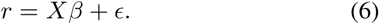

Here *β* is a sparse vector indicating which variables in *X* influences the outcome *r* and 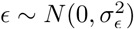.

Note that the components of *r* are uncorrelated—in order to simulate a scenario with correlated observations, we introduce a covariance matrix to simulate correlations between the *y_i_*. Thus, we generate the final response *y* as follows:

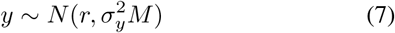

where *M* is the covariance between observations and 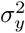 is a scalar that controls the magnitude of the variance. Letting *C* be the matrix formed by stacking the centroids *c_g_*, we choose *M* = *CC^T^*. This has the desired effect of making observations from the same group *g* more correlated.

### B. Experimental Results in Variable Selection

We use the parameters described in Table I in our simulations.

**Table I:**
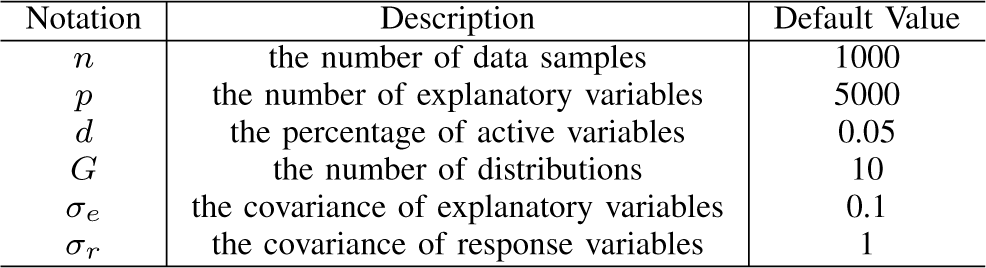
Simulations configurations

The results are shown as ROC curves in Figure 2. In general, across all the parameter settings tested, we see that the proposed Truncated-rank Sparse Linear Mixed Model outperforms the other methods. Unsurprisingly, the Sparse Linear Mixed Model outperforms traditional sparse variable selection methods, which was completely ineffective in this experiment. This illustrates how methods that do not account for possible sources of confounding can drastically underperform when the assumption that observations are independent is violated.

**Figure 2:**
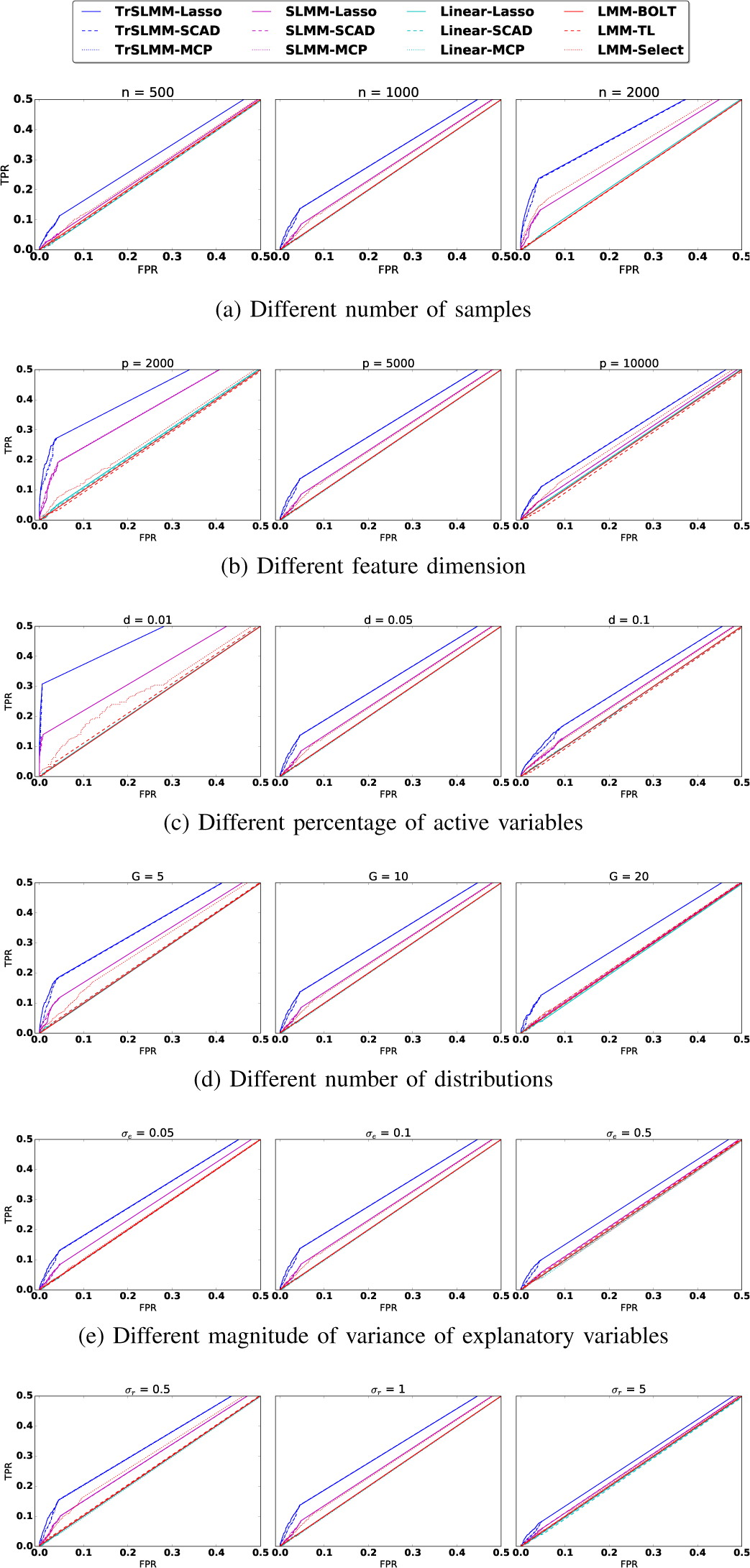
ROC curves for the variable selection experiment. We have zoomed-in to focus on the region of most interest. For each configuration, the reported curve is drawn over five random seeds.

As the various parameters are changed, we observe some expected patterns. For example, in Figure 2(a), as *n* increases, and in Figure 2(b) as *p* decreases, the ratio of 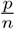 gets smaller and the performance gets better. As we increase the proportion of nonzero coefficients in *β*, the number of distributions, or the variance of response variable *y*, the problem becomes more challenging. In almost all of these cases, however, the TrSLMM-based methods show improved performance. As an example where the SLMM methods are comparable when *G* = 2 SLMM-MCP and SLMM-SCAD behave better than TrSLMM-Lasso, but even they remain slightly inferior to TrSLMM-MCP and TrSLMM-SCAD. Traditional variable selection methods, for the most part, show the same behavior as these parameters are manipulated—this suggests that the fluctuations we observe in the other methods are due to the different strategies by which confounding is corrected.

### C. Prediction of True Effect Sizes

Figure 3 shows the averaged mean squared error in estimating the effect sizes *β* and its standard error over five runs for different settings when we adjust the feature covariance *σ*_*e*_ on synthetic data. We do not consider the LMM extensions here because they do not estimate the effect sizes. Interestingly, we can see that TrSLMM-Lasso behave the best in estimating *β*, while SLMM-Lasso closely follows-up. Traditional sparse variable selection methods (Linear-Lasso, Linear-SCAD, Linear-MCP) behave worse than these two methods, but mostly better than other TrSLMM and SLMM based methods.

**Figure 3:**
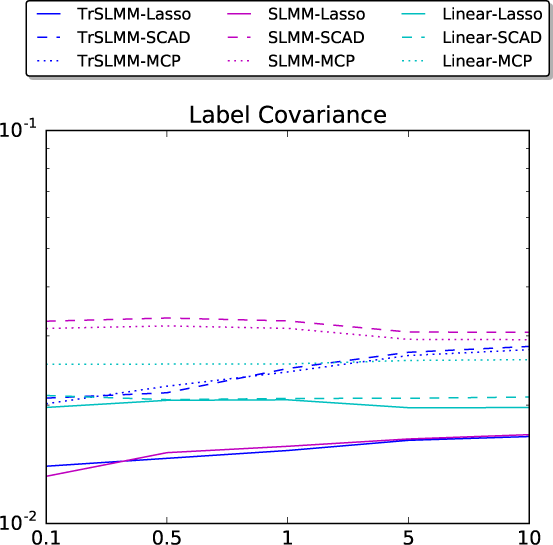
Mean squared error and its standard error with the prediction of true *β*.

### D. Running Time

After confounding correction, we observed that the final sparse variable selection step converged faster. Across all the configurations of synthetic experiments, in comparison to the vanilla sparse variable selection methods, TrSLMM-Lasso, TrSLMM-SCAD, and TrSLMM-MCP only required 49%, 38%, and 29%, respectively, of the time needed for the Lasso, SCAD, and MCP, respectively, to converge on average. SLMM-Lasso, SLMM-SCAD, SLMM-MCP were slightly faster, and only required 28%, 38%, 37% of the time needed on average. While not necessarily faster overall, this is an interesting observation and confirms previous theoretical work suggesting that variable selection is faster and easier for uncorrelated variables.

## V. REAL GENOME DATA EXPERIMENTS

In order to evaluate the TrSLMM framework in a practical setting, we tested our model on three datasets coming from genomics studies. To provide a clearer evaluation, we tested our method on datasets from three different species. We then evaluate our discovered knowledge with some of the published results in relevant literature to show the reliability of our methods compared with existing approaches. Finally, we report our discovered associations. We do not consider the performance of LMM-family models because we have showed their inferior performance in the simulations. Here, we can always attach the truncated-rank idea to these methods and propose new models. We do not believe it is necessary to exhaust these ideas when we can prove the concept of truncated-rank models by comparing vanilla LMM and the truncated-rank counterparts sufficiently.

### A. Data Sets

#### 1 Arabidopsis thaliana

The Arabidopsis thaliana dataset we obtained is a collection of around 200 plants, each with around 215,000 genetic variables [33]. We study the association between these genetic variables and a set of observed traits. These plants were collected from 27 different countries in Europe and Asia, so that geographic origin serves as a potential confounding factor. For example, different sunlight conditions in different regions may affect the observed traits of these plants. We tested the genetic associations between genetic variables with 44 different traits such as *days to germination, days to flowering, lesioning* etc.

##### 2 Heterogeneous Stock Mice

The heterogeneous stock mice dataset contains measurements from around 700 mice, with 100,000 genetic variables [34]. These mice were raised in cages by four generations over a two-year period. In total, the mice come from 85 distinct families. The obvious confounding variable here is genetic inheritance due to family relationships. We studied the association between the genetic variables and a set of 28 response variables that could possibly be affected by inheritance. These 28 response variables fall into three different categories, relating to the glucose level, insulin level and immunity respectively.

##### 3 Human Alzheimer’s Disease

We use the late-onset Alzheimer’s Disease data provided by Harvard Brain Tissue Resource Center and Merck Research Laboratories [35]. It consists of measurements from 540 patients with 500,000 genetic variables. We tested the association between these genetic variables and a binary response corresponding to a patient’s disease status of Alzheimer’s disease.

#### B. Ground Truth for Evaluation

To evaluate the performance of TrSLMM, we compared the results with genetic variables that have been reported in the genetics literature to be associated with the response variables of interest. For Arabidopsis thaliana, we used the validated knowledge of the genetic associations reported in [36]. For heterogeneous stock mice, the validated gold standard genetic variables were collected from the Mouse Genome Informatics database.^1^ For Alzheimer’s disease, we listed the genetic variables identified by one of our proposed model (TrSLMM-MCP) and verified the top genetic variables by searching the relevant literature. Additionally, since the genetic cause of Alzheimer’s disease is still an open research area, we reported the genetic variables we identified for the benefit of domain experts.

#### C. Selected Groups

We first validate the success of our truncated-rank approaches to identify the truly confounding factors from distributions of eigenvalues. Figure 4 shows the distribution of eigenvalues of *XX^T^*. A naïve linear mixed model will correct the confounding factors with all these eigenvalues, resulting in an over-correction. In contrast, Truncated-rank Sparse Linear Mixed Model only identifies the ones that are likely to be confounding sources. As Figure 4 shows, TrSLMM conveniently identifies 27 data origins for Arabidopsis thaliana, while these 200 plants are in fact collected from 27 countries. TrSLMM identifies 65 sources for mice data, while these mice are from 85 different families. Although TrSLMM didn’t pinpoint every confounding factor exactly, the number of confounding factors is much closer compared to vanilla sparse variable selection methods (only one) and vanilla SLMM methods (number of samples by construction). On the human Alzheimer’s Disease, there is no consensus number of data sources available to check the correctness of TrSLMM’s selection, but the distribution seems to indicate that there are only a few confounding sources.

**Figure 4:**
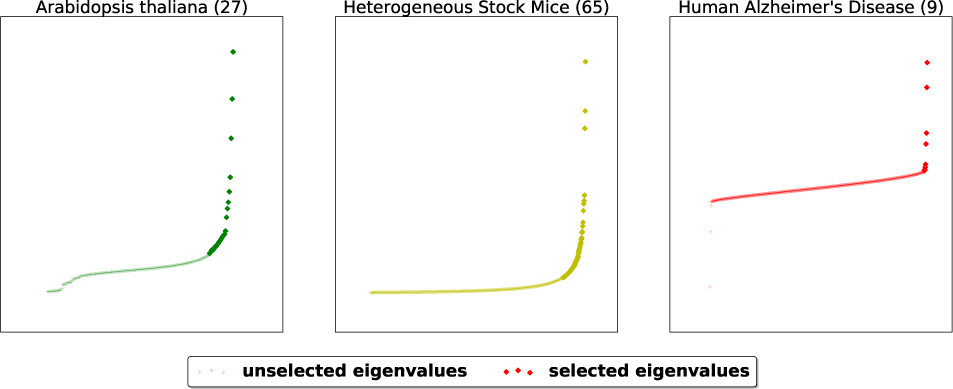
The selected eigenvalues to consider as the sources of confounding factors.

#### D. Numerical Results

Since we have access to a validated gold standard in two out of the three datasets, Figure 5 and Figure 6 illustrate the area under the ROC curve for each response variables (observed trait) for Arabidopsis thaliana and Mice, respectively. The responses are ordered such that the leftmost variables are those for which our TrSLMM model outperform the others. Because discovering associations in genetic datasets is an extremely challenging task, many of these methods fail to discover useful variables. It is worth emphasizing that the discovery of even a few highly associated variants can be significant in practice. Overall, TrSLMM methods managed to outperform the other methods for almost 60% of response variables. TrSLMM-MCP and TrSLMM-SCAD behave similarly, as previously observed in the synthetic data experiments.

**Figure 5:**
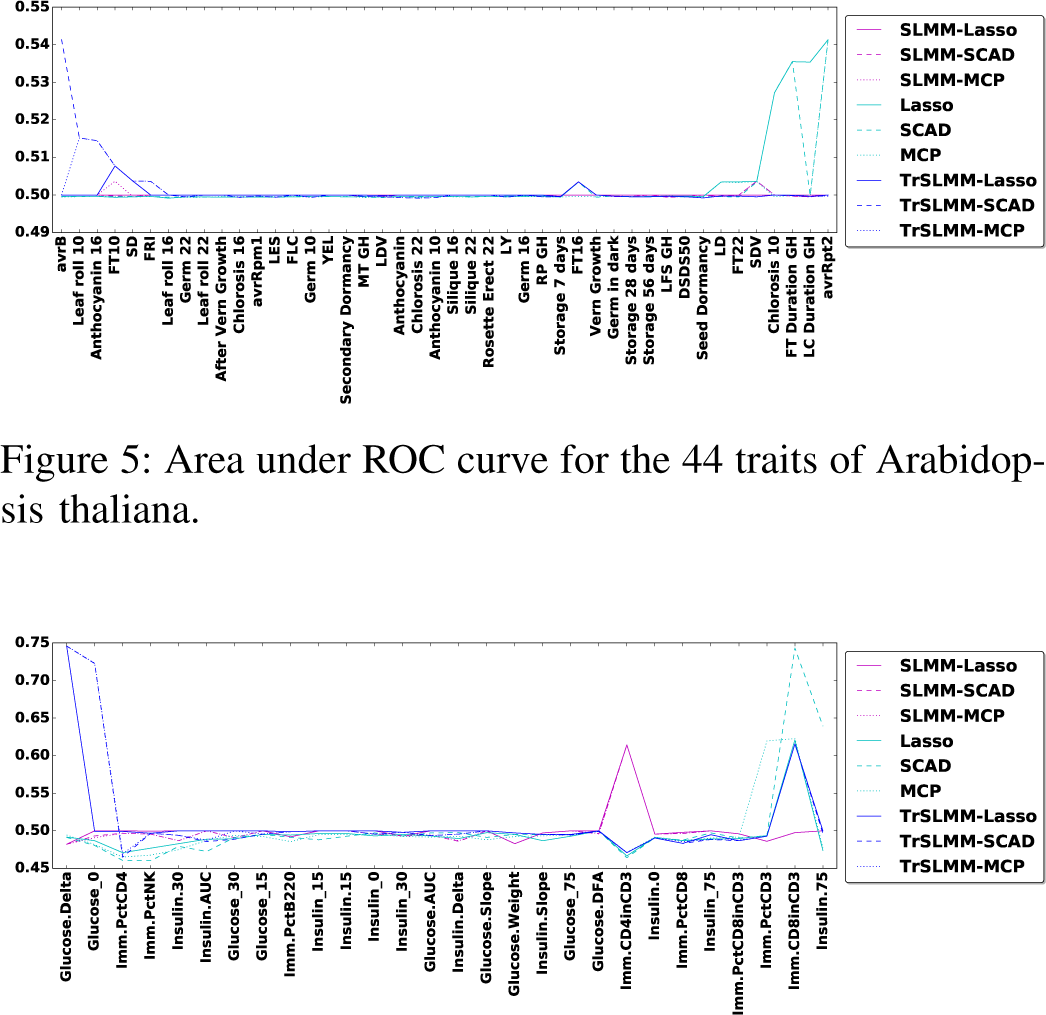
Area under ROC curve for the 44 traits of Arabidopsis thaliana.

**Figure 6:**
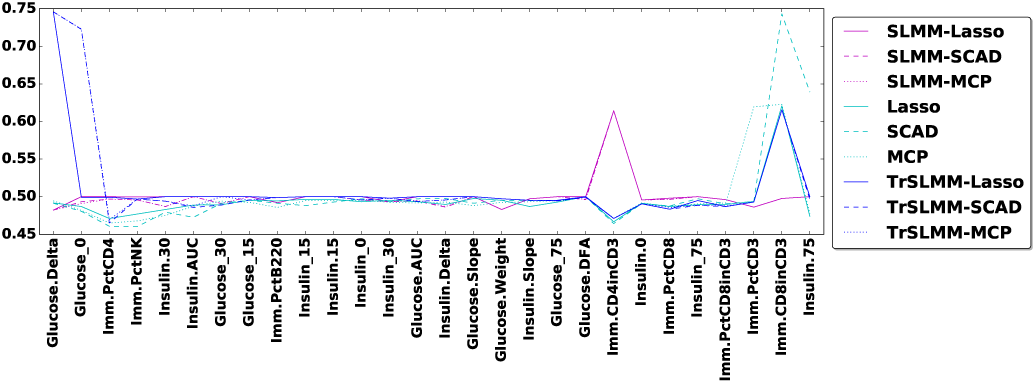
Area under ROC curve for the 28 traits of Mice.

For Arabidopsis thaliana, TrSLMM based models behave as the best one on 56.8% of the traits. Since not all of the traits in our dataset are expected to be confounded, it is not surprising that in some cases traditional methods perform well. Without confounding, one expects methods that are optimized for i.i.d. data to perform best (e.g. Lasso, SCAD, MCP). For example, traits with **GH** in the name mean that the corresponding traits were measured in a greenhouse, where conditions are strictly controlled and potential confounding effects introduced by different regions are minimized. As Figure 5 shows, traditional sparse variable selection methods almost gain the most advantage over greenhouse traits.

For Heterogeneous Stock Mice, TrSLMM based models behave as the best one on 57.4% of the traits. The results are interesting: The left side of the figure mostly consists of traits regarding the amount of glucose and insulin in the mice, while the right hand side of the figure mostly consists of traits related to immunity. This raises the interesting question of whether or not immune levels in stock mice are largely independent of family origin.

Most importantly, our proposed model is at least as good as other SLMM based methods, and sometimes significantly better when confounding is present. This gain in performance comes with no extra parameters and no extra computation, except for one computationally trivial step of screening singular values.

#### E. Knowledge Discovered and Causality Analysis

Finally, we proceed to the Human Alzheimer’s Disease dataset. Because Alzheimer’s Disease has not been studied as extensively as plants and mice, there is no authentic golden standard to evaluate the performances. Here, we report the top 30 genetic variables our model discovered in Table II to foster relevant research.

**Table II:**
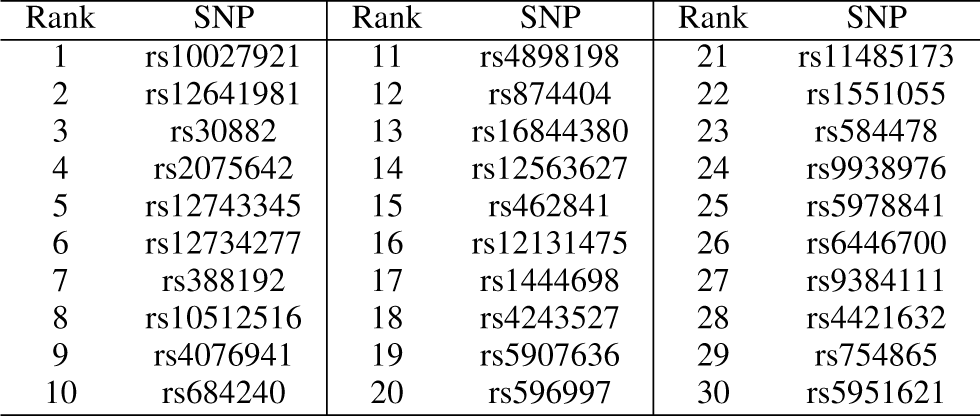
Discovered Genetic Variable with TrSLMM-MCP

| Rank | SNP | Rank | SNP | Rank | SNP |
| --- | --- | --- | --- | --- | --- |
| 1 | rs10027921 | 11 | rs4898198 | 21 | rs11485173 |
| 2 | rs12641981 | 12 | rs874404 | 22 | rs1551055 |
| 3 | rs30882 | 13 | rs16844380 | 23 | rs584478 |
| 4 | rs2075642 | 14 | rs12563627 | 24 | rs9938976 |
| 5 | rs12743345 | 15 | rs462841 | 25 | rs5978841 |
| 6 | rs12734277 | 16 | rs12131475 | 26 | rs6446700 |
| 7 | rs388192 | 17 | rs1444698 | 27 | rs9384111 |
| 8 | rs10512516 | 18 | rs4243527 | 28 | rs4421632 |
| 9 | rs4076941 | 19 | rs5907636 | 29 | rs754865 |
| 10 | rs684240 | 20 | rs596997 | 30 | rs5951621 |

Due to space limitations, we briefly justify only a few of the most important genetic variables here to evaluate the accuracy of our model. The 1^st^ is associated with *ARHGAP10* gene (also called *GRAF2*), which affects the developmentally regulated expression of the *GRAF* proteins that promote lipid droplet clustering and growth, and is enriched at lipid droplet junctions [37]. The 3^rd^ discovered genetic variable is corresponded to *apoB* gene, which can influence serum concentration in Alzheimers disease [38]. The 4^th^ discovered SNP resides within the region of *AOPE*, which is prominently believed to be cause Alzheimer’s disease [39]. The 5^th^ discovered SNP is within *COL1A1*, which is associated with *APOE* [40]. The 6^th^ resides in *WFDC1* and the 9^th^ one is within *GALNTL4*, both are reported to be related with Alzheimer’s disease respectively [41], [42].

## VI. CONCLUSIONS

In this paper, we aim to solve a critical challenge in variable selection when the data is not i.i.d. and does not come from the same distribution. Due to confounding, traditional variable selection procedures tend to select variables that are not relevant. When the sources of confounding are known and can be controlled for, linear mixed models have long been used to make such corrections. The use of LMMs to *implicitly* correct for confounding that is not explicitly known to an analyst is a recent development and a very active area of research. This type of situation occurs frequently in genomics applications where confounding arises due to population stratification, batch effects, and family relationships.

To overcome this problem, we introduced a general framework for sparse variable selection from heterogeneous datasets. The procedure consists of a confounding correction step via linear mixed models followed up by sparse variable selection. We have shown that state-of-the-art variable selection methods such as SCAD and MCP can be easily plugged into this procedure. Further, we showed that the traditional linear mixed model can easily fall into the trap of utilizing too much information, resulting in an over-correction. To correct for this, we introduce a Truncated-rank Sparse Linear Mixed Model that effectively and automatically identifies the sources of confounding factors. Most importantly, we proposed a data-driven, adaptive procedure to automatically identify confounding sources from the spectrum of the kinship matrix without prior knowledge. Through extensive experiments, we exhibited how TrSLMM has a clear advantage over existing methods in synthetic experiments and real genome datasets across three different species: plant (Arabidopsis thaliana), mice, and human.

In future work, we plan to explore more complex structured problems with our proposed framework to select variables for response variables that are dependent [43] or for explanatory variables that are correlated [27]. Further, we plan to integrate our method into the popular genomic research toolbox GenAMap [44].

## ACKNOWLEDGEMENT

This material is based upon work funded and supported by the Department of Defense under Contract No. FA8721-05-C-0003 with Carnegie Mellon University for the operation of the Software Engineering Institute, a federally funded research and development center. This work is also supported by the National Institutes of Health grants R01-GM093156 and P30-DA035778.

## Footnotes

1 http://www.informatics.jax.org/

